# Mapping the Inter- and Intra-genic Codon Usage Landscape in *Homo sapiens*

**DOI:** 10.1101/2025.07.27.667039

**Authors:** Maahil Arshad, Matthew Uchmanowicz, Vanshika Rana, Brett Trost, Stephen W. Scherer, Muhammad Arshad Rafiq

## Abstract

Although the genetic code is degenerate, codon selection is non-random and reflects significant functional constraints. Codon usage bias (CUB) acts as a layer of post-transcriptional regulation, influencing mRNA stability, translation kinetics, and co-translational protein folding. While CUB is well-characterized in unicellular organisms, its regulatory scope and functional consequences in humans remain complex and less defined. Our study offers a comprehensive evaluation of human codon usage. We report that genes exhibiting the strongest codon bias are enriched in high-stoichiometry biological processes, such as skin development and oxygen/carbon dioxide transport, and harbor significantly fewer synonymous variants than expected (ρ = -0.24, *p* < 2.2 x 10^-16^). Furthermore, we find that codon optimization is spatially regulated: it is significantly more pronounced in structured protein domains compared to intrinsically disordered regions (IDRs) (Cliff’s Δ= 0.26, p < 2.2 x 10^-16^). Consistent with translational selection, the most frequently used codons are supported by higher tRNA gene copy numbers (ρ = 0.49, *p* < 6.4 x 10^-4^). Finally, by correcting for GC3 content, we reveal that the apparent correlation between ENC and adaptation indices (CAI/tAI) vanishes, allowing us to disentangle mutational pressure from translational selection. Collectively, our findings position codon usage bias as a central, evolutionarily conserved regulator of translation and protein folding in humans. Our results provide a comprehensive and integrated view of intergenic and intragenic codon usage bias in humans, reinforcing the biological relevance of synonymous codon choice in shaping translational dynamics and protein biogenesis. This provides a refined framework for interpreting synonymous variation and guiding functional genomics.

## Introduction

The redundancy of the genetic code permits multiple codons to specify a single amino acid, a phenomenon known as degeneracy. This redundancy has likely evolved as a buffering mechanism against errors caused by spontaneous and endogenous mutations during DNA replication, transcription, and repair. However, it also provides a regulatory layer that enables differential codon usage, leading to Codon Usage Bias (CUB), the preferential use of certain codons for a given amino acid over others^1^. CUB is driven by several molecular mechanisms, including wobble base pairing at the third codon position, tRNA modifications, codon–tRNA pool correlations, and codon co-occurrence in relation to neighboring codons^2,3^. Consequently, CUB influences numerous processes such as gene expression regulation, mRNA secondary structure formation and the modulation of protein folding and stability.

CUB manifests in two different dimensions: between genes (intergenic), and within individual genes (intragenic). Intergenic bias typically reflects evolutionary pressure for translational efficiency, where highly expressed genes are enriched in optimal codons, leading to increased protein abundance^4^. In contrast, intra-genic bias describes the non-uniform distribution of codon usage within a single transcript, which serves to modulate the local kinetics of the ribosome. Rather than proceeding at a constant rate, translation follows a ’rhythm’ of acceleration and deceleration dictated by tRNA availability^5^. Strategic clusters of rare codons act as ’pausing’ signals, slowing elongation to provide the necessary time for the nascent polypeptide chain to adopt its native conformation co-translationally. Accordingly, structured protein regions tend to use optimal codons to ensure accuracy, while Intrinsically Disordered Regions (IDRs) favor non-optimal ones, reflecting distinct translational requirements^6,7^. Consequently, synonymous variants that disrupt this rhythm can impair protein folding and function, highlighting the non-neutral nature of silent substitutions^8,9^. While the influence of codon usage on translation speed is documented, such as in 5’ ’codon ramps’ that prevent ribosomal congestion, its genome-wide impact on human protein architecture remains under-explored^10,11^. Specifically, the relationship between codon bias and structural disorder is poorly defined. If codon usage actively shapes the folding landscape, we expect limited codon usage in structured domains because a certain translational speed may necessitate correct protein structure. Conversely, IDRs may exhibit diverse codon usage patterns as the lack of secondary structures may allow a variety of codons to be used, consistent with relaxed constraints on translation kinetics since precise ribosomal pausing is not required to guide folding. Crucially, if this restricted codon usage is functional, we predict that these highly biased genes will be selectively constrained against synonymous variation.

Moreover, the metrics traditionally used to assess codon usage bias have inherent limitations, often capturing overlapping signals or being confounded by background nucleotide composition. Consequently, to precisely study CUB in humans necessitates exploring a wide variety of metrics in relation to one another. These include the Effective Number of Codons (ENC), which quantifies the overall magnitude of codon bias by measuring the deviation from random codon usage (where lower values indicate stronger bias)^12^. To estimate translational efficiency, we computed the Codon Adaptation Index (CAI), which evaluates how closely a gene’s codon usage resembles a reference set, such as highly expressed genes, and the tRNA Adaptation Index (tAI), which assesses the compatibility between codon choice and the cellular tRNA pool. Furthermore, we utilized Relative Synonymous Codon Usage (RSCU) to determine specific codon preferences independent of amino acid composition, and GC3 content (GC frequency at the third codon position) to serve as a proxy for background mutational pressure^12–15^. By integrating these diverse metrics, we aim to disentangle genuine translational selection from confounding mutational biases.

To address these gaps, we performed a systematic computational analysis across ∼20,000 human protein-coding genes. Specifically, this study seeks to (1) characterize the genome-wide distribution of key codon-bias metrics in humans, evaluating their inter-correlations to determine the most robust metric for measuring bias; (2) identify specific biological pathways and functional ontologies that are enriched for genes exhibiting either extreme codon usage bias or minimal bias; (3) investigate how codon usage correlates with intrinsic protein structural features (IDRs vs. Structured domains); and (4) assess whether genes with exceptionally restricted codon repertoires are selectively constrained against synonymous mutations.

## Methods

We developed a pipeline integrating high-quality genomic and transcriptomic resources. Canonical coding sequences were curated from Ensembl to calculate codon usage metrics using the cubar R package and custom Python scripts^16^. To assess codon preference, we used tRNA gene copy numbers retrieved from GtRNAdb to calculate tAI, and regional structural disorder annotations were integrated using IUPred3 predictions to differentiate between IDRs and structured domains^17,18^.

Constraint scores from Genome Aggregation Database (gnomAD) were also included to explore the relationship between synonymous codon usage and mutational intolerance^19^. Together, this framework provides a scalable strategy to dissect the biological implications of codon usage across diverse molecular and evolutionary contexts. All statistical analyses were performed in R. P-values < 2.2 x 10^-16^ are reported to indicate probabilities below the floating-point precision limit.

### Datasets

Sequencing data was obtained from Ensembl using release 112, GRCh38.p14 assembly. We retrieved the complete CDS FASTA files from each repository, restricting our analysis to transcript sequences for which a CDS was available. Canonical transcript definitions were retrieved from Ensembl BioMart (release 112), and only transcripts explicitly marked as “canonical” by Ensembl were retained for downstream analyses because they are the most representative of all exons of a gene. This filtering yielded 19 601 high-confidence protein-coding transcripts used throughout the study.

In addition, tRNA genes were obtained from the high-confidence set provided by gtRNAdb v2.0 (downloaded via https://gtrnadb.ucsc.edu/genomes/eukaryota/Hsapi38/hg38-tRNAs.tar.gz), yielding 428 distinct tRNA sequences with their corresponding copy numbers for codon–tRNA pairing analyses.

Gene-level constraint metrics (e.g., synonymous observed vs expected ratio, LOEUF) were obtained from gnomAD v4.1 by downloading gnomad.v4.1.constraint_metrics.tsv from the gnomAD public data release (https://storage.googleapis.com/gcp-public-data--gnomad/release/4.1/constraint/gnomad.v4.1.constraint_metrics.tsv). These metrics were joined to our Ensembl transcript list via stable IDs to compare patterns of synonymous mutation constraint against codon-bias measures.

### Codon-Bias Metric Definitions

All codon usage metrics, including ENC, CAI, tAI, and GC3, were computed using the cubar package (v1.1.0) in R and custom Python scripts. The mathematical definitions, formulas, and interpretive ranges for all metrics are detailed in **Table 1**. RSCU values were calculated for all 19,601 canonical transcripts. To ensure robust global estimates for correlation analyses, we utilized the standard RSCU per codon (CUBAR RSCU). RSCU was also computed across all transcripts after removing outliers, defined as values falling outside 1.5 the Interquartile Range (IQR) of the distribution for that codon (Average RSCU). To distinguish selection-driven bias from background nucleotide composition, ΔENC was calculated by decoupling the observed ENC (*ENC_obs_*) from the expected ENC (*ENC_exp_*) predicted by the gene’s GC3 content (see **Table 1** for Wright’s Curve formulation). A positive ENC (*ENC_exp_ > ENC_obs_*) was used to identify transcripts exhibiting stronger codon usage bias than expected under neutral mutational pressure^20^.

| Metric | Definition | Formula | Range & Interpretation | Source |
| --- | --- | --- | --- | --- |
| **RSCU (Relative Synonymous Codon Usage)** | Ratio of observed codon frequency to expected frequency if all synonymous codons were used equally. | $RSCU_{ij} = \frac{X_{ij}}{\frac{1}{n_i} \sum_{k=1}^{n_i} X_{ik}}$ | Values > 1: Indicates the codon is used more frequently than expected (over-represented). Values < 1: Indicates the codon is used less frequently than expected (under-represented). Value = 1: Indicates random/uniform usage. | Sharp & Li (1987)<sup>13</sup> |
| **ENC (Effective Number of Codons)** | Quantifies the absolute magnitude of codon bias independent of gene length. Measures deviation from random usage. | $N_c = 2 + \frac{9}{F_2} + \frac{1}{F_3} + \frac{5}{F_4} + \frac{3}{F_6}$ | Range is 20 to 61: 20 is Extreme bias (only one codon is used for each amino acid). 61 is No bias (all synonymous codons are used equally). | Wright (1990)<sup>12</sup> |
| **Wright's Curve (Expected ENC)** | Theoretical curve describing the expected ENC value ($ENC_{exp}$) if codon bias were determined solely by GC3 content. | $ENC_{exp} = 2 + GC3 + \frac{29}{GC3^2 + (1 - GC3)^2}$ | Acts as a neutral baseline reference. Points falling on this curve have codon usage explained entirely by nucleotide composition. | Wright (1990)<sup>12</sup> |
| **$\Delta ENC$ (Delta ENC)** | Measure of selection-driven codon bias, calculated by decoupling intrinsic bias from background nucleotide composition. | $\Delta ENC = ENC_{exp} - ENC_{obs}$ | Positive (>0): Bias is stronger than expected from GC3 content (suggests selective pressure). Near or below 0: | Novembre (2002)<sup>20</sup> |
|  |  |  | Bias is largely explained by GC3 content (mutational pressure). |  |
| **CAI (Codon Adaptation Index)** | Measure of translational efficiency based on the resemblance of codon usage to a reference set. | $CAI = \exp\left(\frac{1}{L} \sum_{k=1}^L \ln(w_k)\right)$ | Range is 0 to 1. Value of 1 indicates the gene uses only the most optimal codons found in reference genes' set. While a value of 0 indicates the gene uses the least optimal codons. | Sharp & Li (1987)<sup>13</sup> |
| **tAI (tRNA Adaptation Index)** | Measure of translational efficiency based on genomic tRNA gene copy numbers (tGCN) and wobble pairing rules. | $tAI = \left(\prod_{k=1}^L w_k\right)^{\frac{1}{L}}$ | Range is 0 to 1. Value of 1 indicates codon usage is perfectly optimized to the cellular tRNA pool. While a value of 0 indicates codon usage is poorly matched to tRNA availability. | dos Reis et al. (2004)<sup>14</sup> |
| **GC3 (GC Content at 3rd Position)** | Fraction of Guanine and Cytosine nucleotides at the third codon position; a proxy for mutational pressure. | $GC3 = \frac{Count(G_3) + Count(C_3)}{Total\ Codons}$ | Range is 0 to 1. Value of 1 indicates that all codons end in G or C (complete GC bias). While a value of 0 indicates that all codons end in A or T (complete AT bias). | Muto & Osawa (1987)<sup>21</sup> |

### Programs

Ensembl protein-coding FASTA sequences were filtered via a custom R-based quality control pipeline to ensure that every sequence represented a complete open reading frame (ORF). Specifically, we verified that sequences initiated with a canonical AUG start codon, terminated with one of the three standard stop codons (UAA, UAG, UGA), and contained no premature termination codons. Sequences failing these criteria were discarded prior to downstream codon usage analyses. The remaining sequences were then loaded into R using Biostrings, and headers were parsed to extract the transcript ID, Ensembl gene ID, gene symbol, and GRCh38 genomic coordinates. These metadata were compiled into a unified table linking each sequence to its genomic context and used in all subsequent cubar analyses. FASTA entry name was reset to the transcript ID to ensure a one-to-one link between sequence and metadata. Next, triplet frequencies were tabulated for every transcript against the standard nuclear codon table. The same counts were subsequently used to derive all downstream metrics. Finally, all metrics were merged with the transcript-level metadata to yield a final table containing transcript ID, gene ID, gene symbol, genomic coordinates, CAI, tAI, ENC and GC3s. This file represents a comprehensive codon-bias atlas for the human proteome and serves as the basis for all subsequent comparative analyses.

Coding DNA sequences (CDS) for all human protein-coding transcripts from GRCh38 (Ensembl release 112) served as the starting dataset. Each FASTA header was parsed to extract transcript (ENST) and gene (ENSG) identifiers; only the first instance of every unique ENST-ENSG pair was kept to eliminate redundancy. CDSs whose lengths were not divisible by three were removed, and the remaining sequences were translated in silico with the standard genetic code, truncating at the first in-frame stop codon. Proteins shorter than ten amino acids were discarded, and every surviving sequence was saved to an individual FASTA file named “ENST_ENSG.fa”.

These protein FASTA files were analysed locally with IUPred3 (v3.2) in long mode, which assigns a disorder probability (0–1) to each residue. Residues scoring ≥ 0.50 were labelled intrinsically disordered (IDR), whereas those < 0.50 were considered structured.

For every transcript, nucleotide coordinates of IDR residues and of structured residues were gathered and concatenated into two separate sequences: one representing all disordered segments and the other all structured segments. Together they formed a composite FASTA containing exactly two entries per transcript. All downstream codon-usage analyses were performed in R 4.4.1 with the cubar package, exactly as described previously, to calculate codon bias metrics for IDRs and structured regions for each transcript.

Additionally, categorization of genes into three distinct categories was based on the distribution of their ENC and ΔENC values. Thresholds were established using specific quantiles calculated from the full dataset. Translational Selection genes were defined as those in the bottom 20th percentile of ENC (indicating high bias) that simultaneously fell in the top 20th percentile of ΔENC (indicating bias deviating from GC3 expectations). Mutational Bias genes were defined as those in the bottom 20th percentile of ENC but with a ΔENC value below the 80th percentile. Unbiased Control genes were defined as those in the top 20th percentile of ENC values.

Functional enrichment analysis was performed using the clusterProfiler package in R. Gene symbols were mapped to Entrez IDs using the org.Hs.eg.db annotation database. We performed Gene Ontology (GO) enrichment for Biological Processes (ont = “BP”) using the enrichGO function. The gene universe (background) was defined as the complete set of genes analyzed in the codon usage metrics. Significance was assessed using a False Discovery Rate (FDR) adjustment (pAdjustMethod = “fdr”) with a significance cutoff of *q* < 0.05. For visualization, the top 15 significantly enriched terms (sorted by FDR) were plotted.

Analyses were performed using the following packages (versions): cubar (1.1.0), Biostrings (2.72.1), data.table (1.16.2), dplyr (1.1.4), tidyverse (2.0.0), ggplot2 (3.5.1), ggpubr (0.6.0), GGally (2.2.1), patchwork (1.3.0), gghalves (0.1.4), ggtext (0.1.2), glue (1.8.0), rstatix (0.7.2), ppcor (1.1), and knitr (1.48). Supplementary processing was performed in Python used biopython (1.85) and pandas (2.2.3). All scripts are available at github.com/maahilarshad/CodonBias.

## Results

As established by the 1000 Genomes Project, a typical diploid human genome varies from the reference genome at about 4.1 to 5 million sites, majority of which are heterozygous Single Nucleotide Polymorphisms (SNPs)^22^. However, since protein-coding regions account for only ∼1% of the genome (approx. 30 Mb), coding variation is relatively sparse. On average, SNPs occur every 600–1,000 nucleotides, implying a total of 30,000–50,000 coding SNPs genome-wide.

While an average gene may harbor 2–3 SNPs, this count naturally correlates with gene length. Crucially, each individual carries only ∼9,600 synonymous variants on average, suggesting that any single gene is likely to harbor one or fewer synonymous variants per individual^23^. Consequently, codon-usage profiles are unlikely to vary significantly between individuals, supporting the validity of calculating metrics from the reference genome. We chose protein-coding sequences from Ensembl to study codon usage metrics across the human genome, and all subsequent correlation analyses are Spearman-based (all metrics are non-normally distributed; Fig. S1-S2).

### Distribution of Codon Usage Metrics

Across ∼20,000 Ensembl protein-coding genes (Fig. 1A), ENC values (blue) range from 33.01 to 57.73 (mean = 49.35; median of 50.58; IQR = 46.31– 52.95), reflecting moderate gene-to-gene variability in overall codon bias. GC3 content (gold) spans the widest interval of 0.16 to 1.00 (mean = 0.59; median = 0.59; IQR = 0.44–0.73), indicating substantial heterogeneity in third-position GC composition. In contrast, CAI (green) is tightly clustered between 0.55 and 0.96 (mean ≈ median = 0.80; IQR = 0.77–0.83), while tAI (orange) ranges from 0.20 to 0.45 (mean ≈ median = 0.29; IQR = 0.27–0.31), both showing comparatively constrained variation in codon-adaptation indices. The lower absolute values and narrow spread of tAI likely reflect incomplete annotation of human tRNAs: because tAI is calculated from the set of codon-cognate tRNAs in reference, any uncharacterized or low-abundance isoacceptors are omitted, leading to an underestimation of the true translational adaptation.

**Figure 1.**
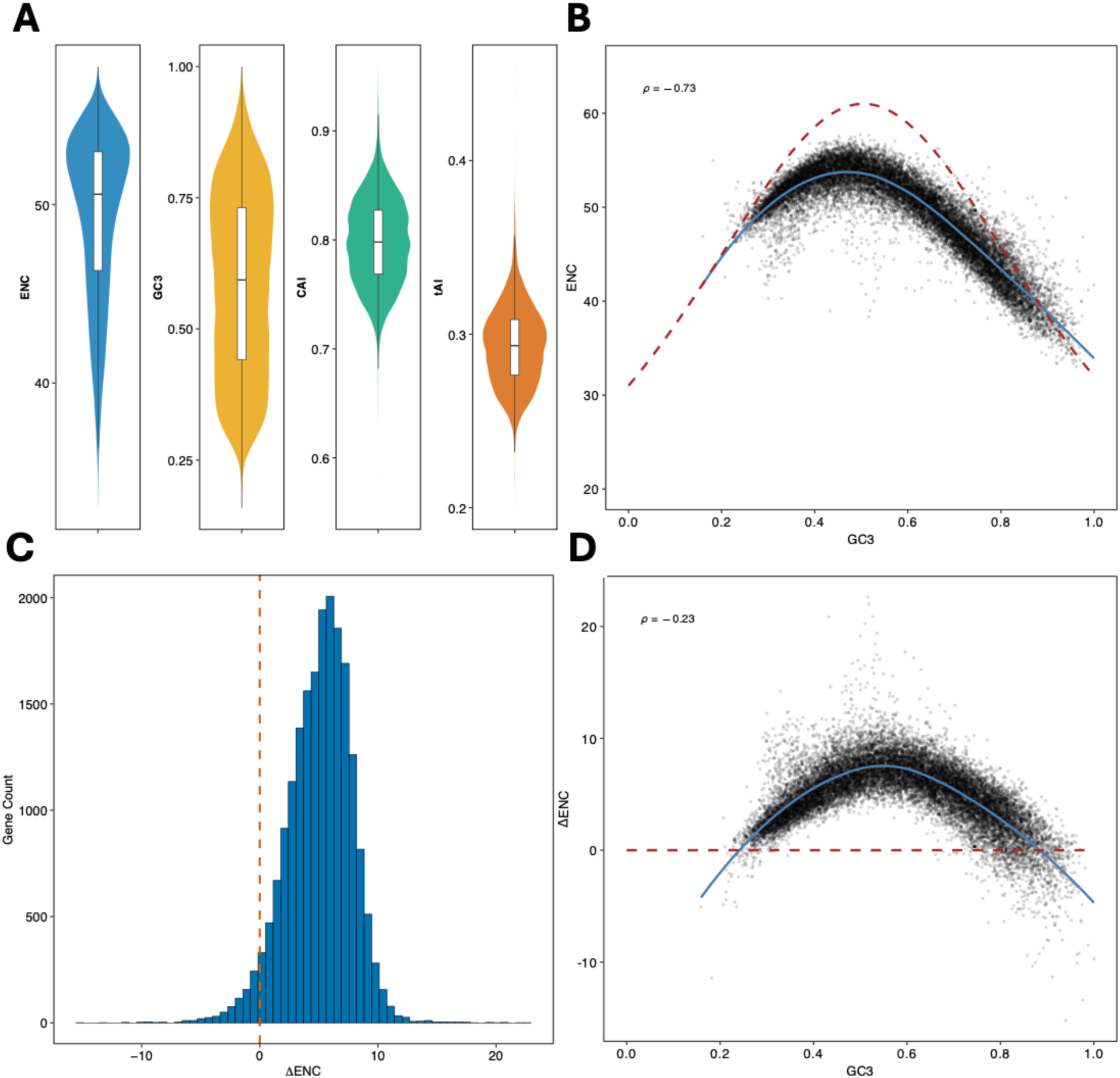
Codon-bias landscapes across the human Ensembl protein-coding gene set. **(A)** Violin plots with overlaid boxplots showing the genome-wide distributions of ENC, GC3, CAI, and !fil_for ∼20,000 human protein-coding genes. **(B)** Scatterplot of GC3 versus ENC for every protein-coding gene, overlaid with Wright’s neutral expectation cuNe (red dashed) and a LOESS smooth curve (solid blue cuNe; Spearman p = -0.73). **(C)** Histogram showing the distribution of ΔENC (ENC corrected for GC3 content). The vertical orange dashed line (at 0) indicates the neutral expectation, highlighting a shift toward positive values. **(D)** Scatterplot of GC3 versus ΔENC for every gene, overlaid with the neutral expectation baseline (red dashed line) and a LOESS smooth curve (solid blue cuNe; Spearman *p* = -0.23).

Figure 1B plots GC3 versus ENC for all genes and overlays Wright’s neutral-expectation curve (red dashed line). Most points fall below the neutral expectation (red dashed line), a trend tracked closely by the empirical LOESS fit (blue solid curve) indicating stronger codon bias than GC3 alone predicts. A Spearman rank correlation revealed a strong negative relationship (ρ = –0.73), while the LOESS trend (blue curve) highlights the non-linear nature of this decline: as GC3 increases, ENC decreases. Consequently, low ENC values can arise in part from elevated GC3 rather than from genuine translational selection, underscoring the need to correct for nucleotide composition when identifying genes with codon-usage bias independent of GC3. Therefore, to detect genes biased independent of GC3 content, we used a metric called ΔENC, which can be calculated by considering expected ENC based on GC3 content (as per Wright’s neutral-expectation curve) minus the observed ENC calculated directly from the sequence^20^. The distribution of ΔENC ranges from –15.23 to 22.63 (median = 5.3; mean = 5.04; IQR = 3.35–6.91); because positive ΔENC values indicate stronger codon bias than predicted by GC3 alone, these summary statistics show that most genes exhibit a modest excess of codon bias, whereas only a small subset (the negative tail) is less biased than expected (Fig. 1C). Interestingly, plotting GC3 against ΔENC, Wright’s neutral curve collapses to the ΔENC = 0 baseline because the GC3-driven component of codon bias has been subtracted out (Fig. 1D). Most genes nevertheless lie above this red-dashed line, indicating additional bias beyond nucleotide composition, and the residual correlation is weakened as compared to the use of ENC (ρ = –0.23). This attenuation shows that GC3 now explains little of the remaining variation, making ΔENC a cleaner, composition-independent metric that more precisely captures true codon-usage bias.

### Functional Enrichment of Differentially Biased Genes

Codon usage lies on a spectrum, making any single threshold for classifying bias arbitrary. Moreover, the non-normal distribution of codon bias metrics necessitates the use of quantiles to assess genes at the extremes of this spectrum. Therefore, gene categories were defined using specific quantile thresholds for both ENC and ΔENC (Fig. 2A).

**Figure 2.**
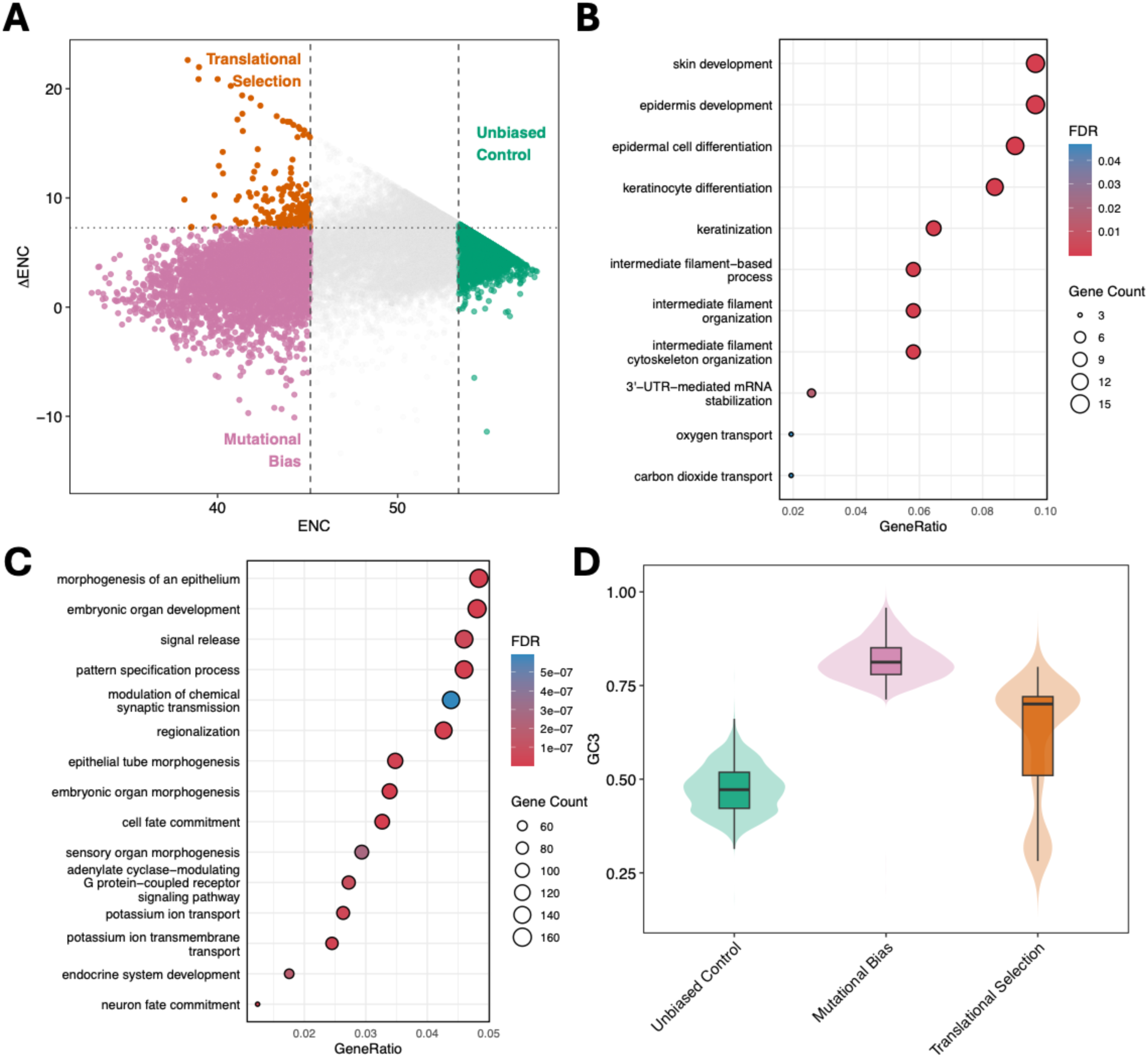
Functional enrichment and compositional analysis of gene sets defined by codon usage patterns. **(A)** Scatterplot of ENC versus *t::,.* ENC. Genes are classified into three distinct categories based on their deviation from neutral expectations: Translationally Selected (orange; low ENC, high t::,.ENC), Mutationally Biased (pink; low ENC, low t::,.ENC, and Unbiased Controls (green; high ENC). **(B)** Dot plot showing the top enriched Gene Ontology (GO) biological processes for the Translationally Selected genes. **(C)** Top enriched GO terms for the Mutationally Biased genes. For Band C, dot size represents gene count and color indicates significance (FDR). **(D)** Violin plots with overlaid boxplots comparing the distribution of GC3 content across the three gene categories.

The vertical dashed lines mark the ENC thresholds, while the horizontal dashed line marks the ΔENC threshold. Translational Selection genes (orange) are defined as the 20% of genes with the lowest ENC (left of the first vertical line) that also possess a ΔENC in the top 20^th^ percentile (above the horizontal dashed line); these genes maintain bias despite GC3 correction. Mutational Bias genes (pink) are 20% with the lowest ENC but with a ΔENC below the 80^th^ percentile (below the horizontal line), indicating their bias is largely driven by GC content. Unbiased Control genes (green) are 20% of genes with the highest ENC values (right of the second vertical line), which cluster near a ΔENC of 0.

Next, a GO biological processes enrichment test identified pathways enriched in each category. Genes subject to translational selection were enriched in processes involving structural integrity and transport, specifically skin/epidermis development, keratinization, and oxygen/carbon dioxide transport (Fig. 2B). Genes characterized by mutational bias were enriched in developmental and signaling pathways, including epithelial morphogenesis, synaptic transmission, and neuron fate commitment (Fig. 2C). Finally, unbiased genes appear to be involved in regulatory processes affecting non-coding RNAs—perhaps genes that do not require rapid production in large amounts, with enrichments in ncRNA processing, regulatory ncRNA-mediated gene silencing, and piRNA processing (Fig. S3E).

Finally, an analysis of GC3 content reveals distinct distributions among the three categories (Fig. 2D). While mutationally biased genes exhibit high GC3 content (median ∼0.8) and unbiased control genes show intermediate GC3 content (median ∼0.5), genes undergoing translational selection display a wide range of GC3 values. This broad distribution illustrates that translational selection operates independently of specific GC3 content.

### Relationships between Codon-Bias Metrics

As codon–usage bias intensifies (lower ENC), genes gravitate toward optimal codons that are decoded by abundant tRNAs, so both CAI (*ρ =* –0.64) and tAI (*ρ =* –0.63) increase (Fig 3A). This inversely proportional relationship is expected because CAI is explicitly designed to reward codons that appear more often in a given set of sequences, while tAI weights codons by the genomic copy-number of their cognate tRNAs, a quantity still imperfectly measured in human due to extensive modifications and tissue-specific expression of tRNAs, lowering absolute tAI values^24^.

**Figure 3.**
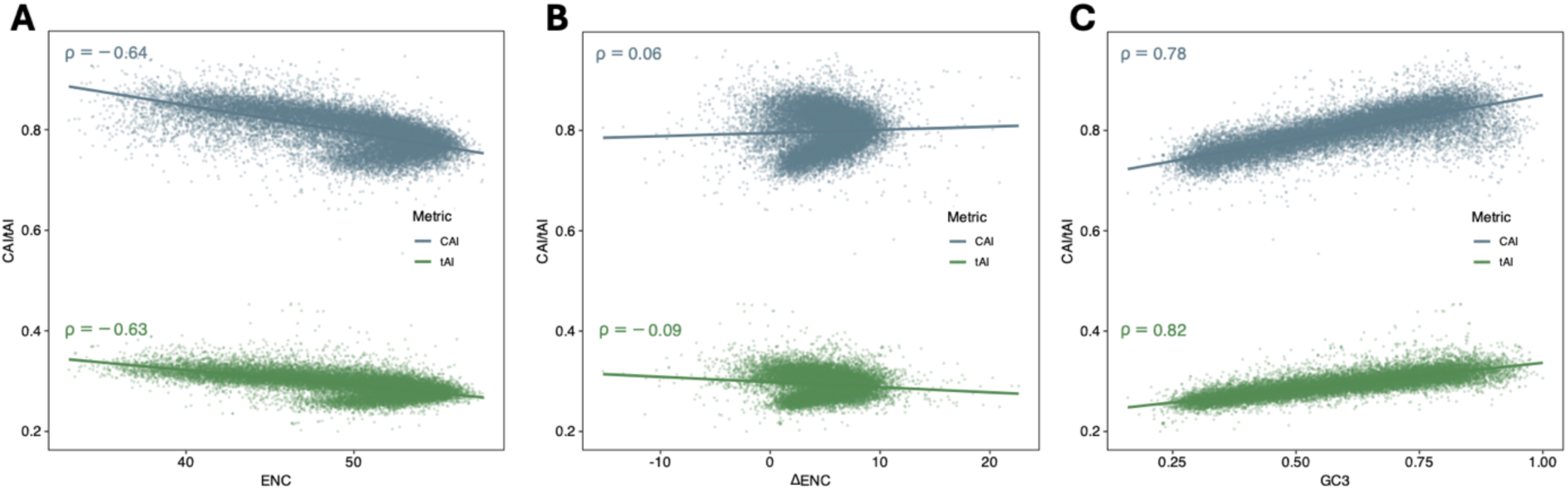
Assessing the effect of GC3, ENC, and ΔENC on translational efficiency metrics CAI and tAI. Gene•level scatterplots of codon adaptation index (CAI, blue) and tRNA adaptation index (1M, green) against three codon-bias measures: (A) ENC versus CAI <u>and tAI showing</u> strong negative correlations (CAI: *p* = --0.64; *r* = - 0.63), indicating that genes with fewer used codons exhibit higher translational adaptation. (B) ΔENC versus adaptation indices, revealing minimal association (CAI: *r* = 0.06;tAi *r* = -0.09), which implies that selection on codon usage beyond GC3 content has little impact on these metrics. (C) GC3versus CAI and tAI with strongpositive correlations (CAI:*r* =0. 78; *r* = 0.82), demonstrating that nucleotide composition is the primarydriver of codon·adaptation variation.

When we correct ENC for its GC3 dependence (ΔENC), these links almost vanish (CAI: *ρ =* 0.06; tAI: *ρ =* –0.09; Fig 3B). In other words, once base-composition bias is removed, there is little residual signal of extra translational selection captured by either CAI or tAI. This interpretation is reinforced by Fig 3C: GC3 alone explains the bulk of variation in both CAI (*ρ =* 0.78) and tAI (*ρ =* 0.82), mirroring earlier genome-wide reports that CAI and tAI track GC-rich codons^25^.

ENC’s theoretically expected relationship with GC3 comes from Wright’s neutral curve, which allows us to compute ΔENC and disentangle true codon-usage bias from mere nucleotide composition. No comparable neutrality model exists for CAI or tAI: CAI is intrinsically tied to high-expression or set of sequences chosen, and tAI inherits any compositional skew in the tRNA gene repertoire, both factors make it difficult to generate a GC3-independent expectation for these metrics. Consequently, downstream analyses will treat ΔENC as the primary GC3-corrected measure of codon bias, while CAI and tAI will be interpreted cautiously, acknowledging that their apparent adaptation largely reflects underlying GC3 enrichment rather than additional translational-selection pressure.

### Intrinsically Disordered Regions and Codon-Usage Bias

Multiple studies have reported that structured protein regions exhibit stronger codon-usage bias than IDRs in various species, leading to the hypothesis that optimal codons may facilitate co-translational folding and stable tertiary structure^6,26^. To test this genome-wide in humans, we retrieved all human protein-coding transcripts from Ensembl and ran IUPred3 to score per-residue disorder (cutoff ≥0.5 to define IDRs). Within each transcript, we concatenated all IDR segments and all structured segments separately, then computed key codon-usage metrics (ENC, ΔENC, CAI, tAI and GC3) for each region class. This region-level design allowed a direct comparison of codon bias between disordered and ordered segments across the proteome (Fig. 4). After doing so, we found that IDRs illustrate a higher median ENC, thus weaker codon-usage bias than structured segments. In our dataset, IDRs have a median ENC of 52.79 (mean 52.69; IQR = 50.29–55.40), whereas structured regions show a median ENC of 50.65 (mean 49.44; IQR 46.50–52.90). This shift is significant and small-to-moderate in magnitude (Cliff’s Δ = 0.33; 95 % CI = 0.32–0.34; p < 2.2 × 10⁻¹⁶; Fig. 4A). After correcting for GC3 content, IDRs show a lower median ΔENC of 2.85 (mean 0.53; IQR = –1.92 – 5.96) compared with structured regions (median 5.02; mean 4.45; IQR 2.79–6.83), with a small-to-moderate effect size (Cliff’s Δ = 0.26; 95 % CI = 0.25–0.27; p < 2.2 × 10⁻¹⁶; Fig. 4B), confirming that codon usage is more biased in structured regions and is not simply driven by nucleotide composition. By contrast, while CAI, tAI and GC3 each reached nominal significance by Wilcoxon rank-sum (p < 2.2 × 10⁻¹⁶), the actual differences are relatively small once effect size is considered: Cliff’s Δ = 0.07 for CAI, 0.06 for tAI, and 0.05 for GC3 (all considered negligible by Cliff’s criteria; Fig. 4C, S3A, S3B, S3C). This trend where even trivial shifts become significant in very large samples, confirms that translational-adaptation indices and base composition are effectively indistinguishable between IDRs and structured regions.

**Figure 4.**
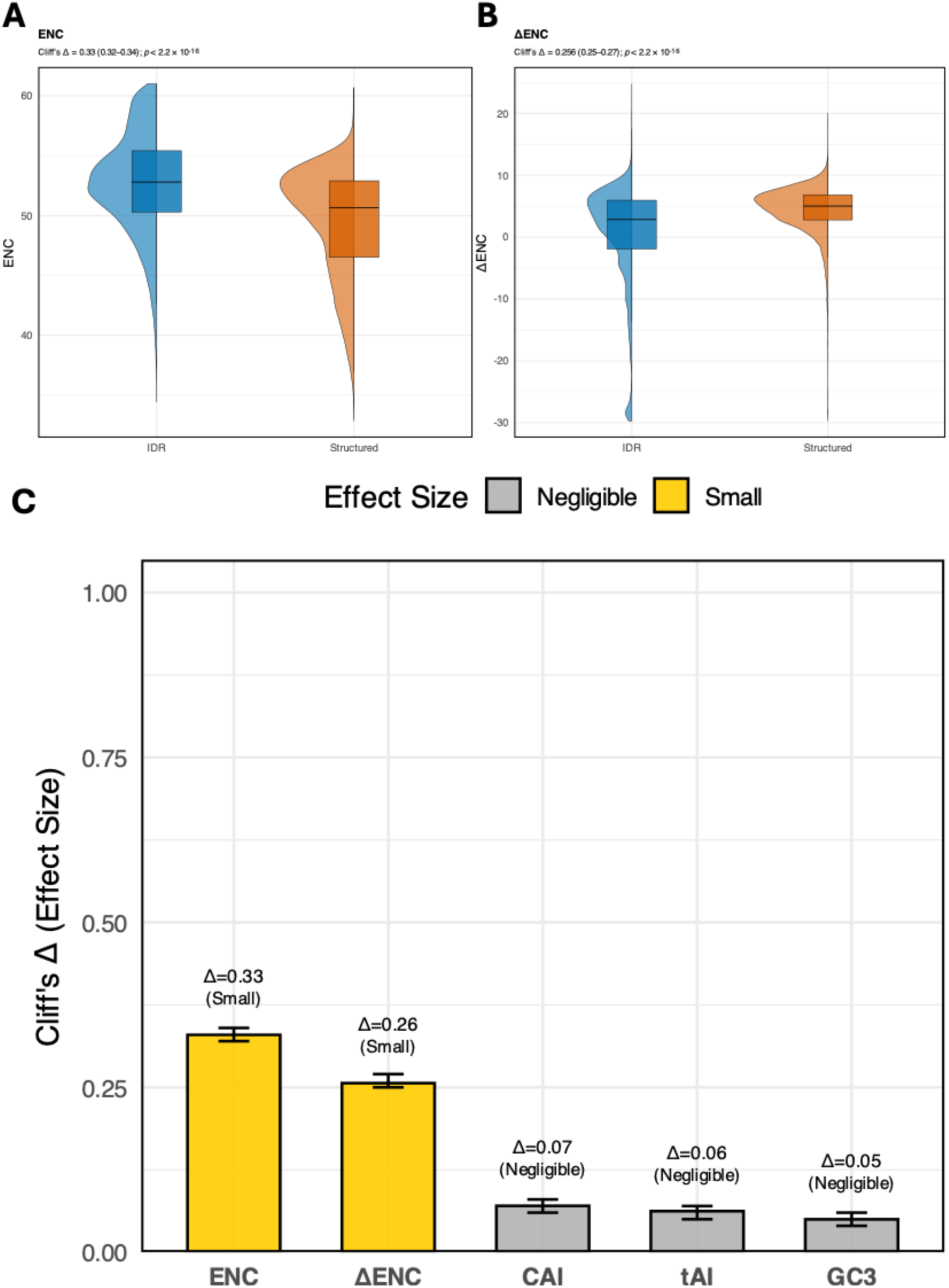
Codon-usage differences between IDRs and structured regions. **(A)** Violin/box plots of ENC show that IDRs (blue) have significantly higher ENC, thus, weaker codon usage bias than structured regions(orange). Statistical comparison by Cliff’s Δ,.yields Δ.= 0.33 (95% Cl 0.32-0.34), *p* < 2.2e-16. **(B)** Violin plotsof t:,.ENC revealthatIDRs exhibit significantly lower Δ,ENC than structured regions (Cliff’s Δ,.= 0.26, 95% Cl 0.25 - 0.27; *p* < 2.2e-16), demonstrating that even after accounting for GC3-driven bias, IDRs maintain weaker codon-usage bias than structured regions. **(C)** Bar chart of Cliff’s *t,.* effect sizes (with 95% confidence intervals) for five metrics comparing IDRs to structured regions: ENC (Δ=0.33, small), ti.ENC (Δ=0.26, small), CAI (Δ=0.07, negligible), tAI (Δ=0.06, negligible) and GC3 (Δ=0.05, negligible). Smalleffects are shown in yellow; negligible effects in gray.

### Linking Codon Usage Bias to Mutation Spectra, tRNA Availability & Stop-Codon Frequencies

Across ∼20,000 human protein-coding genes we explored how codon-usage bias, background mutation pressure, tRNA supply and termination choice intertwine (Fig. 5A–D). ΔENC was negatively associated with the ratio of observed-to-expected synonymous variants (ρ = –0.24, p < 2.2 × 10⁻¹⁶; Fig. 5A). Thus, the more a gene departs from neutral codon usage, the more its preferred codons appear to be protected from silent changes, consistent with pervasive purifying selection at synonymous sites reported in recent genome-wide analyses^27^. By contrast, the per-site synonymous mutation rate (µ) rose only marginally with ΔENC (ρ = 0.02, p = 0.017; Fig. 5B), indicating that mutational rate alone cannot account for the depletion of silent variants in highly biased genes. Similarly, genes with stronger codon bias tend to have lower synonymous constraint scores (LOEUF), suggesting they are less tolerant to synonymous variants than genes with diverse codon usage (Fig. S3D).

**Figure 5.**
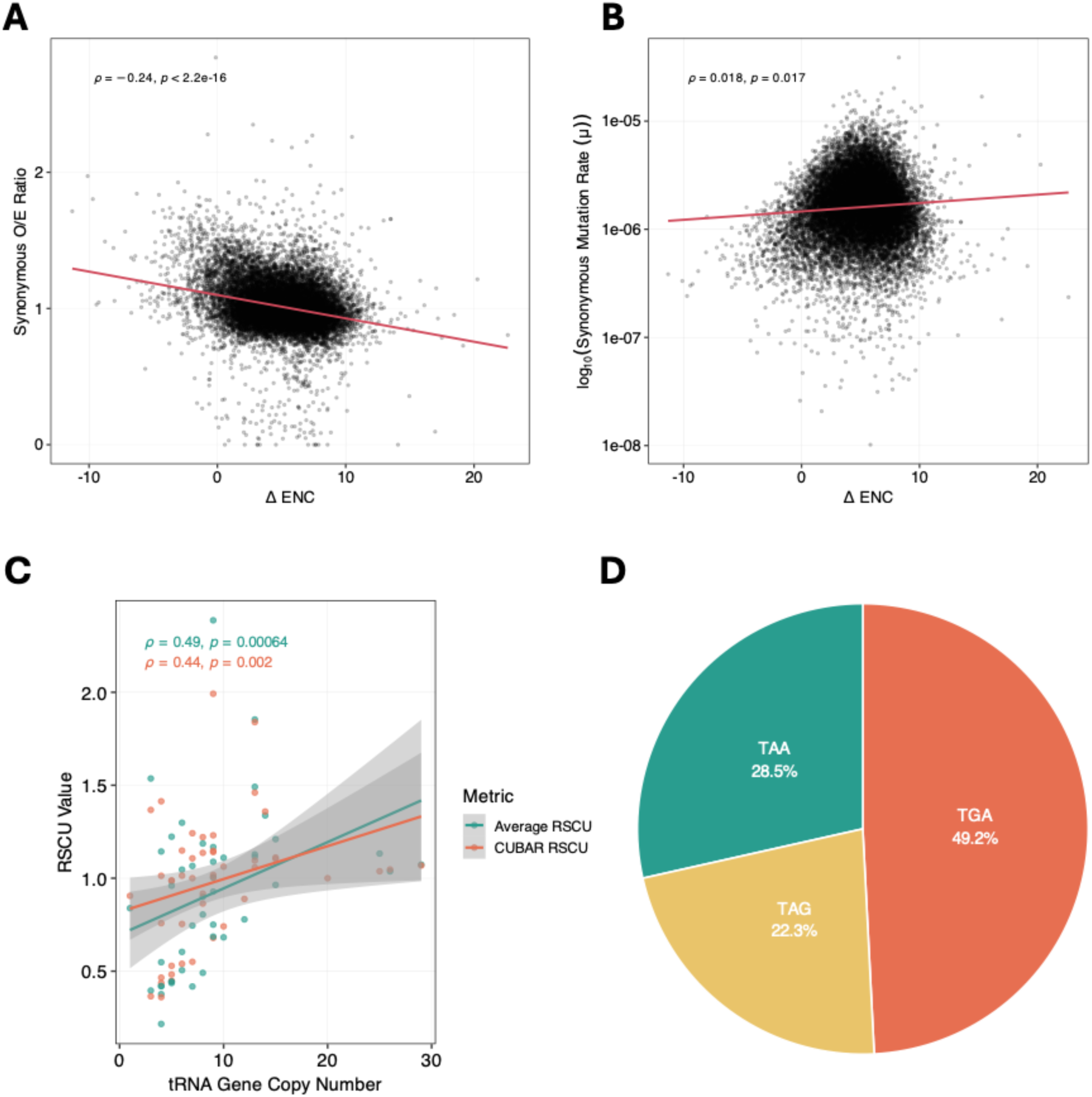
Codon-bias deviation, mutation patterns, tRNA abundance, and stop-codon usage. **(A)** Scatterplot of 6ENC versus the synonymous observed/expected (O/E) ratio. A linear regression fit (red line) shows a significant negative correlation *(p* = -0.24, p < 2.2×10·^16^), indicating that genes under stronger codon bias (higher 6ENC) have fewer observed synonymous variants than expected. **(B)** Scatterplot of 6ENC versus the per-site synonymous mutation rate. A linear regression fit (red line) shows a weak but significant positive correlation (p= 0.02, p < 2.2×10·^16^), indicating that codon-biased genes exhibit slightly higher underlying mutation rates. **(C)** Scatterplot of RSCU per codon against cognate tRNA gene copy number. The plot compares RSCU values calculated for each gene, averaged and outliers removed (teal; *p* = 0.49, p = 0.00064) and the CUBAR RSCUwas calculated genome-wide (orange; *p* = 0.44, p = 0.002). Both sets reveal a moderate positive correlation, showing that codons decoded by more abundant tRNAs are used more frequently. Shaded areas represent 95% confidence intervals for the regression lines. **(D)**Pie chart of genome-wide (∼20,000 protein-codinggenes) stop-codon frequencies: TGAis most common (49.2%), followed by TAA(28.5%) and TAG(22.3%).

We next asked whether codon usage tracks the genomic supply of cognate tRNAs. Averaging RSCU for each codon across each gene, removing outlier genes and plotting it against the copy number of its matching tRNA gene revealed a positive correlation (ρ = 0.49, p = 0.000064; Fig. 5C). Similarly, standard RSCU calculation (CUBAR RSCU) using all genes/codons in the genome maintains the correlation but is slightly lower than the previous method (ρ = 0.44, p = 0.002; Fig. 5C). This pattern mirrors classical codon–tRNA co-adaptation models, where selection favours codons translated by abundant tRNAs to maximise translational efficiency^7^. Finally, genome-wide stop-codon frequencies were markedly skewed: TGA accounted for almost half of all termination codons (49.2%), whereas TAA (28.5%) and TAG (22.3%) were less common (Fig. 5D), consistent with previously reported literature^28,29^. Collectively, these observations highlight that codon bias, mutation constraints, tRNA availability and stop-codon choice are tightly coupled elements of translational optimisation in the human genome.

## Discussion

A growing body of experimental and theoretical work demonstrates that synonymous codon usage is not merely a passive consequence of mutational biases but can actively influence co-translational protein folding. Early experimental evidence showed that silent mutations can alter folding pathways and functional outcomes without changing amino-acid sequence^8^. Subsequent studies proposed that locally rare codons can introduce translational pauses that facilitate the stepwise folding of nascent polypeptides, while coordinated patterns of codon usage may tune the kinetics of translation to match folding requirements^5,30^. These findings suggest that regions of proteins subject to strong structural constraints may experience selective pressure on codon usage to ensure proper folding, whereas intrinsically disordered regions, which lack stable tertiary structure, may tolerate more relaxed codon usage patterns.

Non-random codon usage is a pervasive phenomenon observed across the tree of life and was among the earliest discoveries in molecular biology. However, elucidating its functional roles, such as in protein folding, mRNA stability, and translational efficiency, has historically been challenged by technological constraints. Over the years, as technology improved, it became easier to correlate changes in protein structure to diseases. Any changes that spared protein structure, would inevitably spare protein function, hence coining such mutations as “silent”, earning them lesser importance than mutations that alter protein structure, hence function^31^. Nevertheless, the 1000 Genomes Project revealed that most variation in humans tends to be found in non-coding regions and most plausibly affects gene regulation rather than protein structure^22^. Thus, the quantity and appropriate cellular distribution of proteins are equally important for protein function. This builds a compelling case that any genetic change that alters gene regulation can either drive disease or contribute to phenotypic diversity within a species. Likewise, we argue that codon usage is a means to regulate genes, and genetic changes that alter codon usage have the potential to dysregulate biological function.

Codon usage can be seen as a continuous measure because the extent to which a gene uses codons may reflect the specificity in translational dynamics, that do not need to be the same for all proteins to perform their respective functions. Ribosomal proteins and housekeeping genes or genes constitutively expressed, frequently display codon preferences aligned with abundant tRNAs, supporting stable and efficient protein synthesis. For example, the gene encoding Elongation Factor Tu (EF-Tu) in bacteria exhibits near-perfect codon optimization. Research by Brandis and Hughes, demonstrated that experimentally replacing these optimal codons with synonymous, slower variants caused a significant drop in organismal fitness^32^. This decline occurred because the ’slow’ codons caused ribosomes to linger longer on the mRNA, sequestering the cellular pool of ribosomes and reducing the overall capacity for global protein synthesis. This example illustrates how codon usage bias may encode regulatory information that may help fine tune expression of different proteins, reinforcing its role as a layer of translational control. The most reliable method to study codon usage is ΔENC, which is derived from decoupling ENC from GC3 content^20^. We can see that majority of the genes depict a positive ΔENC value, indicating that they are more biased or limited in their codon usage than that would be expected from GC3 content alone. The ΔENC calculation is inherently absolute, as it relies directly on the exact codon frequencies observed in each transcript minus the effect from GC3 content. Similarly, this trend is also evident from the ENC values, as most genes fall below Wright’s curve (Fig. 1B). Furthermore, employing ΔENC mitigates the collinearity often seen between tAI/CAI and raw ENC values. This distinction is important because correlated metrics obscure the nuance between different evolutionary pressures; if two metrics mirror one another, they do not add explanatory power. ΔENC resolves this by accounting for compositional constraints, making it a superior metric for isolating selection-driven bias in future comprehensive analyses.

Conversely, tAI can be computed in multiple ways but depends on comprehensive data, such as tissue-specific tRNA expression levels or the precise tRNA gene copy number across the genome. When such information is incomplete, as it is in our study, tAI scores may be underestimated and thus offer limited insight into true codon bias. Additionally, this data scarcity precludes the application of the ’correction for isoacceptors’ suggested by Sabi and Tuller, which optimizes wobble penalties (*s*-values) specifically for the target organism; without this correction, the metric relies on fixed, universal weights that likely misrepresent the actual decoding efficiency of the local tRNA pool and should therefore be prioritized in future studies^33^. Likewise, CAI can be derived either from a set of highly expressed genes or from overall codon frequencies in the transcriptome, with the most frequent codons deemed optimal. Although we adopted the transcriptome-wide approach, CAI remains tightly coupled to GC3 content; without an explicit neutrality model like Wright’s curve for ENC, it’s hard to disentangle mutational bias from selection. As such, CAI adds little beyond what GC3 alone reveals, and the same limitation would apply even if we had complete tRNA copy-number data. Yet, we observed a clear trend between tRNA gene copy number and the average RSCU value: as the cognate tRNA copy number increases for a given codon, its RSCU value also increases. Biologically, this is expected, since more frequently used codon tend to be supported by a larger pool of matching tRNAs.

Codon usage bias is inherently continuous, often rendering static thresholds arbitrary and potentially misleading. Furthermore, the non-normal distribution of codon bias metrics in the human genome complicates the use of parametric statistics. To address this, we employed a quantile-based stratification strategy (Fig. 2A). By defining categories relative to the population distribution rather than fixed values, we successfully isolated genes at the true extremes of the spectrum. Crucially, the inclusion of ΔENC as a secondary filter allowed us to disentangle two distinct evolutionary forces: Translational Selection (bias maintained despite compositional constraints) and Mutational Bias (bias driven principally by GC content). This reveals that translational selection in the human genome is not a universal pressure, but a specialized mechanism reserved for “high demand” systems. The enrichment of Translational Selection genes in processes like skin development, keratinization and gas transport are highly significant because these pathways represent the physiological extremes of protein abundance. Epidermal cells and erythrocytes are essentially protein factories requiring the massive, constitutive production of structural (keratins) and cellular scaffolds (cytoskeletal filaments), and transporters (hemoglobins). A critical insight from this study is the characterization of Mutational Bias genes, particularly those enriched in synaptic and neurodevelopmental pathways. If one relied solely on standard ENC values, these genes would appear highly optimized; however, the ΔENC correction reveals that their “bias” is mostly due to the effect of high GC content. This distinction is biologically coherent with the isochore structure of the human genome. Many brain-specific genes reside in GC-rich isochores, which naturally push codon usage toward G/C-ending codons regardless of translational necessity^34^. Unlike the “factory-like” requirements of skin or blood cells, neurodevelopmental processes often rely on precise spatiotemporal timing and complex regulatory networks rather than raw protein abundance. Therefore, the evolution of these sequences appears driven by the local genomic landscape (isochores and methylation contexts) rather than a push for translation speed. This underscores the danger of interpreting low ENC as evidence of selection without accounting for the background nucleotide composition. Finally, the enrichment of Unbiased Control genes in regulatory pathways, such as ncRNA processing and gene silencing, offers an intriguing counterpoint to the high-demand genes. These proteins often function catalytically or stoichiometrically, meaning they are needed in precise, often low, amounts rather than in bulk. For these regulatory genes, a lack of codon bias (random usage) may functionally represent a “neutral” or even beneficial state. Rapid translation is not a priority; in fact, slower, non-optimized translation rates can be advantageous for complex regulatory proteins, allowing sufficient time for co-translational folding or the assembly of multi-subunit complexes.

Beyond the inter-genic variation observed between different gene classes, our results highlight distinct intra-genic patterns driven by protein architecture. Previous reports that codon usage is more biased in structured protein domains than IDRs hints at an interplay between codon usage and protein folding^35^. The more selective usage of codons in structured domains may be necessary for accurate protein folding, while IDRs may use various codons as certain protein regions may not need to assume a permanent structure. IDRs being less structurally constrained than folded domains, frequently exhibit reduced codon bias, consistent with relaxed translational selection. Such regions may therefore serve as evolutionary ‘flexibility zones,’ enabling sequence variation that contributes to intra-species diversity without compromising essential function, as seen in proteins like p53, which combines a structured DNA-binding domain with highly disordered regulatory termini^36^. Given that p53 and other cancer-associated proteins often contain extensive IDRs, our findings support the prediction that codon bias may play an underappreciated role in cancer biology by influencing the regulation of such structurally heterogeneous proteins. Consistent with our results, we see that structured protein regions have a greater median ΔENC suggesting that the codons used in these regions may aid with protein folding, as compared to IDRs that use a greater variety of codons. Although the effect size is small, it may nonetheless reflect underlying biology.

Our study grouped all structured and IDR regions into two broad meta-segments. Yet both structured domains and IDRs comprise multiple subclasses, each of which may exhibit distinct codon-usage patterns^7^. Thus, future work should investigate codon usage across the various classes of structured and disordered regions. More rigidly structured segments will preferentially employ optimal codons as natural selection promotes efficient and accurate translation especially in regions critical for folding and function^37,38^. This specific focus on structural domains directly addresses the gap we highlighted in the introduction: ‘While the influence of codon usage on translation speed is documented, such as in 5’ ’codon ramps’ that prevent ribosomal congestion, its genome-wide impact on human protein architecture remains under-explored.’ Consistently, our method prioritized this architectural link by concatenating segments based on disorder rather than genomic position. A consequence of this design, however, is that position-specific phenomena like 5’ ramps are averaged into the broader meta-segments. Therefore, while we successfully established the link between codon bias and protein structure, future work should employ sliding-window analyses to superimpose positional data onto these structural maps, fully disentangling ’ramps’ from domain-specific constraints.

Next, we predicted that genes with the most constrained codon usage would exhibit fewer observed synonymous variants than expected, especially if codon choice is functionally important. Indeed, we observed that as ΔENC increases (indicating more restricted codon usage), the number of reported synonymous variants falls below expectation. These genes also show greater constraint against synonymous mutations, reflected by lower synonymous LOEUF scores for these genes. In contrast, repeating this analysis with raw ENC values yields the opposite pattern, largely because ENC is influenced by GC3 content, and GC-rich regions are known to have higher mutation rates^39^. Moreover, synonymous mutation rates account for very little of the variance in ΔENC. Despite similar background synonymous mutation rates, genes with more constrained codon usage accumulate fewer synonymous changes, strongly suggesting their functional importance. Future work should therefore investigate synonymous variants in these genes across different disorders. That being said, not every synonymous change in these genes will have an effect: the local codon context and surrounding sequence likely modulate any functional impact. Nonetheless, genes with high ΔENC are prime genome-wide candidates for prioritizing studies that link synonymous mutations to disease.

Finally, the predominance of TGA is well documented in mammalian genomes and is thought to reflect a combination of GC-rich context, differential read-through rates and historical substitution biases^28^. The predominance of TGA as a termination codon in our analysis aligns with previous findings that show TGA usage is enriched in newly evolved genes, possibly due to reduced translational termination signals on the complementary strand and selection-driven context effects that favor TGA over TAA or TAG^29^.

A critical limitation of our broad genomic survey lies in the simplified treatment of the ∼20,000 human genes as equivalent, independent entities. This approach is necessary for initial genome-wide analysis but does not fully account for the complex architectural nature of the human genome, which is shaped by extensive paralogues, gene families, segmental duplications, and retro-transposition events.

The most significant constraint of this uniform approach is its failure to stratify genes by their evolutionary history. Studies have demonstrated a significant association between codon usage and the evolutionary age of genes in metazoan genomes, including humans, suggesting that factors beyond immediate selection pressure shape codon preference^40^. Specifically, by analyzing genes in aggregate, we may obscure the significant impact of gene age and duplication timing on codon usage analysis; for example, recently duplicated genes (paralogues) may not have had sufficient evolutionary time to diverge in their codon preferences compared to their ancestors.

This raises the question of whether ancient, conserved genes (singletons) lacking paralogues, which often represent core cellular machinery, might be underrepresented in current results. These single-copy, essential genes (often called “singletons” or ohnologs) are known to be under significantly higher evolutionary constraint compared to genes from segmental duplications^41^. Their constrained nature suggests that changes to their optimal codon usage would have a disproportionately large functional impact, yet their limited number means their extreme codon bias may be statistically dominated by the signals of larger, evolutionarily younger gene families. Future analyses must therefore stratify genes by their evolutionary history to disentangle the constraints acting on ancient singletons versus rapidly expanding gene families, ensuring that the unique translational dynamics of core cellular components are not overlooked.

## Conclusion

In conclusion, codon usage is a fundamental feature of the human genome, critically shaping a wide range of biological processes. Our analysis reveals that codon optimization is not uniform but spatially and functionally regulated. The pronounced codon bias in structured protein domains, contrasted with the reduced bias in IDRs, suggests a dual evolutionary strategy: structured domains may be constrained to ensure accurate co-translational folding, while IDRs may serve as evolutionary ‘flexibility zones,’ tolerating sequence variation that fosters intraspecies diversity without compromising protein stability. Furthermore, the strong correlation between frequently used codons and tRNA gene copy number points to a co-evolutionary mechanism driven by translational selection, ensuring efficient protein synthesis for high-demand genes.

Crucially, we find that genes exhibiting the strongest codon bias are not random but are enriched in processes requiring massive protein stoichiometry, such as skin development, gas transport, and cytoskeletal organization. The fact that these genes harbor significantly fewer synonymous mutations than expected confirms that codon choice is under strong selective constraint. By correcting for GC3 content, we further demonstrate that this bias is distinct from background mutational pressure, highlighting the functional importance of synonymous sites.

Ultimately, our findings add to a growing appreciation that synonymous changes are not merely silent passengers in the genome but elements of a nuanced genetic language. The choice of codon, much like selecting the precise word in a sentence, encodes regulatory signals that influence translation dynamics and protein expression even when the amino acid sequence remains unchanged. In this view, codon usage acts as a layer of biological communication, embedding regulatory intent into the protein’s genetic blueprint. These insights emphasize the need to revisit the so-called ‘silent’ portion of the genetic code—not as neutral noise, but as a subtle dialect shaping phenotypic diversity, disease susceptibility, and evolutionary innovation.

---

**Figure S1.**
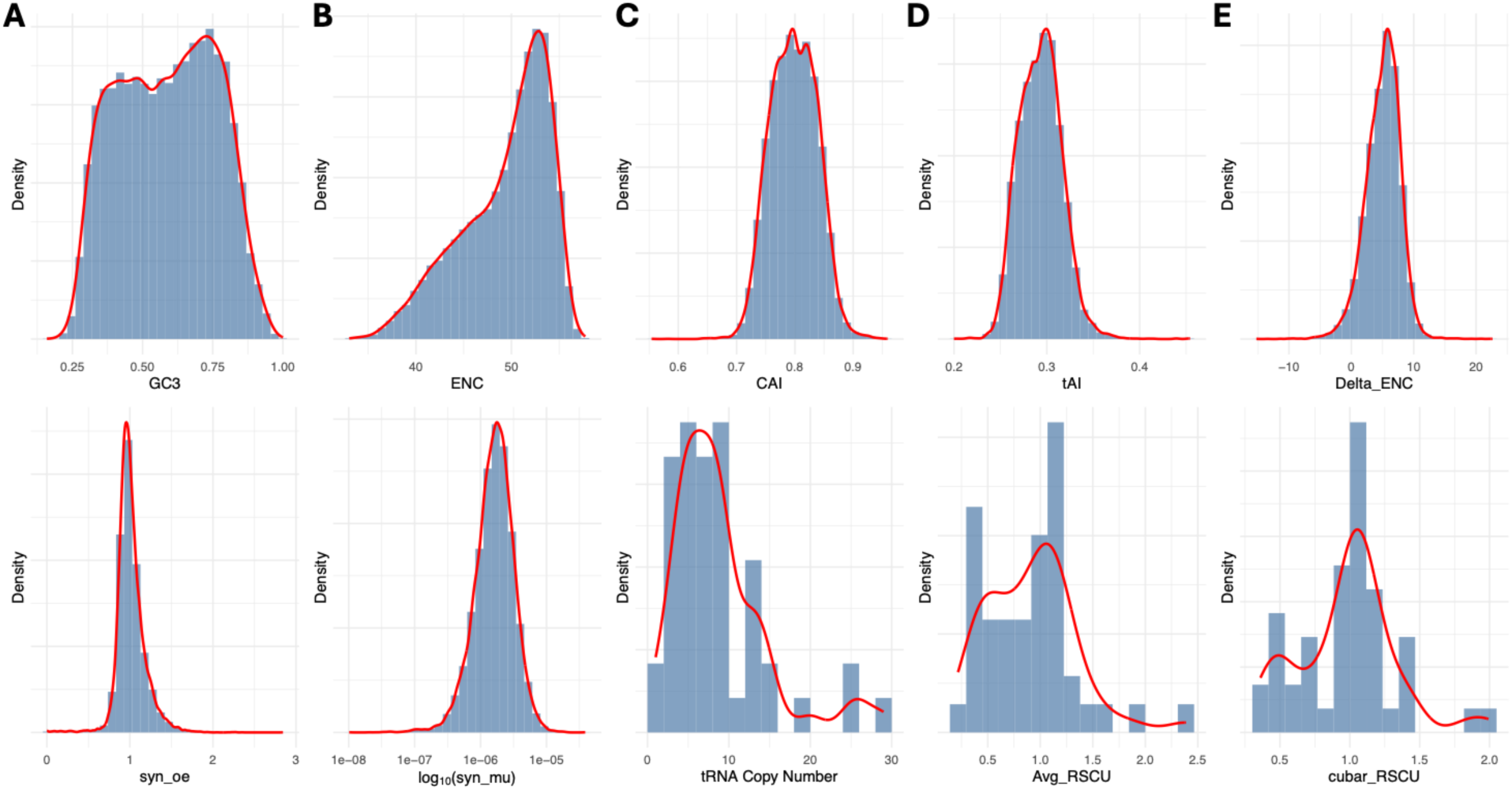
Genome-wide distributions of codon usage metrics. Histograms with overlaid kernel density estimates (red lines) showing the data structure for the variables analyzed in this study. **Top Row**: (**A**) GC content at the third synonymous position (GC3); (**B**) Effective Number of Codons (ENC); (**C**) Codon Adaptation Index (CAI); (**D**) tRNA Adaptation Index (tAI; (E) ΔENC (deviation of ENC from neutral expectation). **Bottom Row**: (F) Synonymous observed/expected ratio <u>(syn. Oe),</u> acting as a proxy for selective constraint: (**G**) Synonymous mutation rate log10 scale: (**H**) tRNA gene copy number: (**I**) Average Relative Synonymous Codon Usage (RSCU) calculated per gene/outlier genes removed; (**J**) CU BAR RSCU calculated genome-wide using CUBAR.

**Figure S2.**
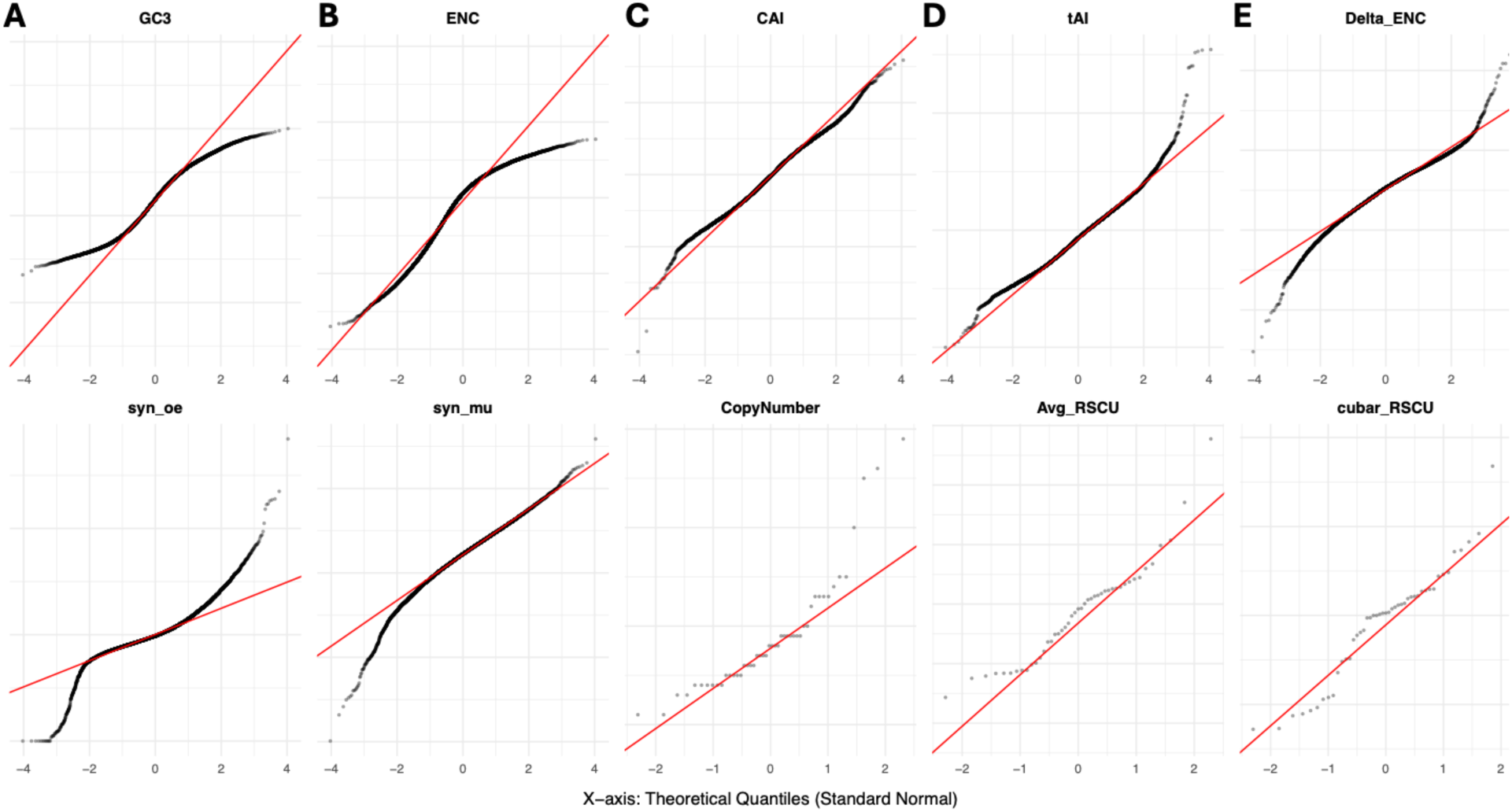
Quantile-Quantile (Q·Q) plots assessing the normality of genome-wlde variables. Each panel displays a Q-Q plot comparing the distribution of the observed data against a theoretical standard normal distribution. TheX-a.xis represents the theoretical quantiles, and the Y-axis represents the observed sample quantiles for each metric. The red solid line serves as a reference for perfect normality; points falling along this line indicate a normal distribution. **Top Row**: (**A**) GC content at the third synonymous position (GC3); (**B**) Effective Number of Codons (ENC); (C) Codon Adaptation Index (CAI); (**D**) tRNA Adaptation Index(CAI; (**D**). ΔENC (deviation of ENC from neutral expectation). **Bottom Row**: (**F**) Synonymous observed/expected ratio (syn oe); acting as a proxy for selective constraint; (**G**) Synonymous mutation rate log1O scale; (H) tRNA gene copy number; (**I**) Average Relative Synonymous Codon Usage (RSCU) calculated per gene/outlier genes removed: (**J**) CU BAR RSCU calculated genome-wide using CUBAR.

**Figure S3.**
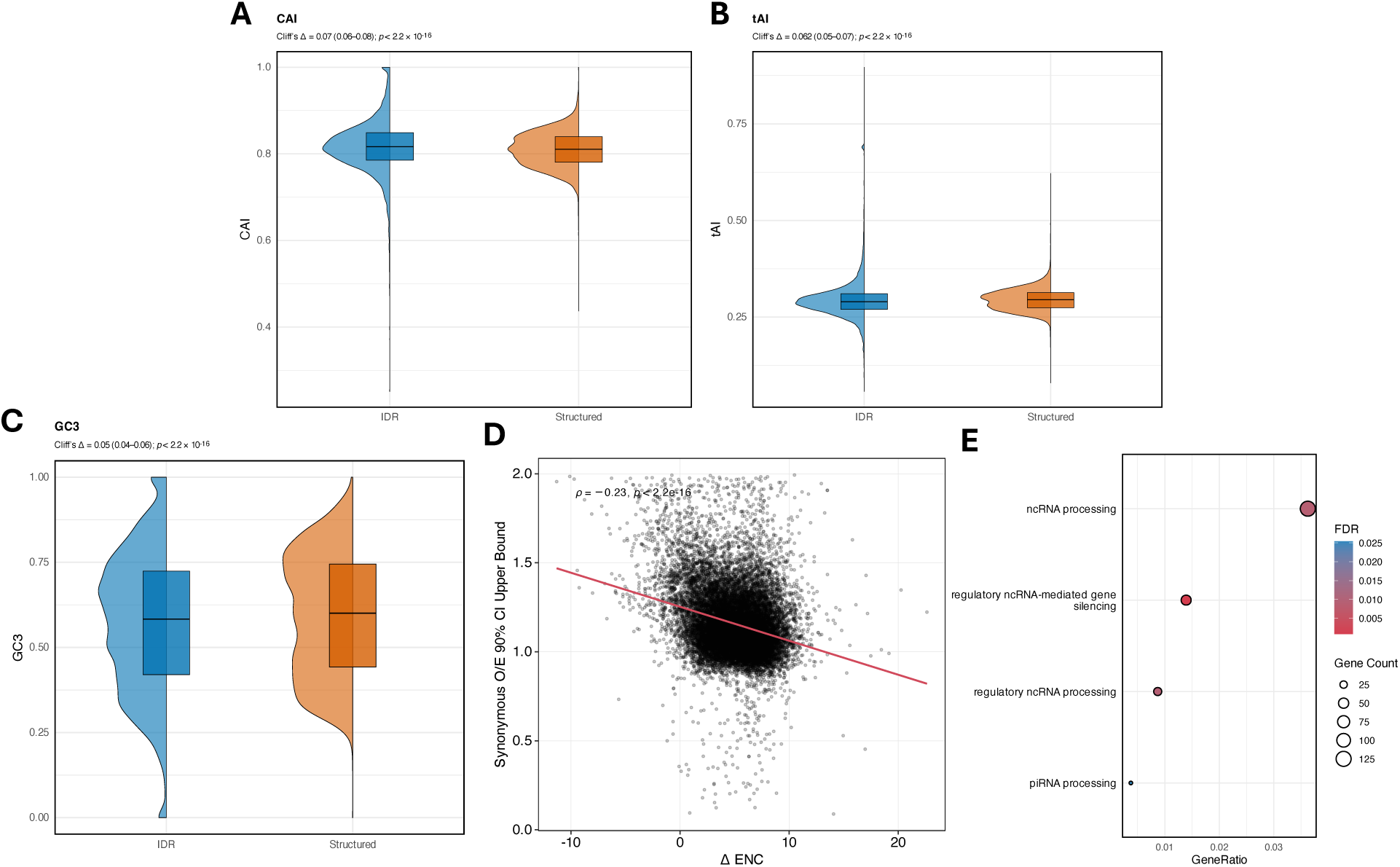
(**A**) Violin and box plots of CAI in IDRs (blue) versus structured regions (orange). Median CAI values are nearly identical, and although the difference is statistically significant, the effect size is negligible (p < 2.2 × 10⁻¹⁶; Cliff’s Δ = 0.07). **(B)** Violin and box plots of tAI show similar distributions in IDRs and structured regions (p < 2.2 × 10⁻¹⁶; Cliff’s Δ = 0.06). **(C)** Violin and box plots of GC3 reveal comparable GC3 content between IDRs and structured regions (p < 2.2 × 10⁻¹⁶; Cliff’s Δ = 0.05). **(D)** Scatterplot of ΔENC versus the upper bound of the 90% CI for synonymous O/E shows a negative correlation (ρ = –0.23; p < 2.2 × 10⁻¹⁶), indicating that genes with stronger codon bias tend to have lower synonymous constraint scores. (**E)** Top enriched GO biological process terms for unbiased control genes. Dot size represents gene count and color indicates significance (FDR).

